# The role of animal personality in behavioural welfare diagnostics: a case study in Arctic charr (*Salvelinus alpinus*)

**DOI:** 10.64898/2026.08.25.747029

**Authors:** Joris Philip, Gabrielle Ladurée, Audrey Prat, Marion Dellinger, Sabine Lobligeois, David Benhaïm

## Abstract

Animal personality is the expression of consistent among-individual variation in a behavioural trait across time and context. The application of this theory to behavioural research provides a valuable framework to investigate the among- and within-individual variation in behaviour. The animal personality theory is particularly relevant to understand and diagnostic fish welfare. It can be integrated within the nature-based welfare framework, which emphasizes the expression of species-specific behaviours and the maintenance of consistent behavioural patterns over time. Because behavioural trait such as boldness may be closely linked to other functional phenotypes, such associations reflect the broader concept of animal personality, whereby a behavioural trait can covary with another phenotypic trait to support an adaptive responses to environmental conditions. Although these relationships are both species- and context-dependent, they are consistently shaped by environmental conditions and environmental complexity tend to promote species-specific behaviours and reduce maladaptive traits. Here we examined how structural environmental complexity shapes personality in Arctic charr and their covarying functional phenotypes, specifically growth rate and brain size. We propose that environmental complexity promotes alternative behavioural and functional phenotypes through multivariate phenotypic plasticity. We found that environmental complexity did not influence mean boldness between both treatments, but repeatability of boldness in the complex environment was remarkably consistent over a longer-term period, while estimates collapse after seven days in the plain treatment. Our findings are a clear evidence that environmental complexity foster stable behavioural trait expression and that a plain environment may suppress personality. Our results provide compelling evidence that behavioural structure and dynamics are embedded within patterns of behavioural variance. Although we found no support for behavioural covariation or associations with growth rate and brain size, we suggest that the animal personality framework may offer a valuable approach for diagnosing fish welfare issues through the partitioning of behavioural variance.

## Introduction

Animal personality is defined as the expression of consistent among-individual variation in a behavioural trait across time and contexts (Dochtermann et al., 2014; Oers & Mueller, 2010; Wilson, 2018). As part of the quantitative genetic framework, the theory of animal personality provides valuable insight into behavioural dynamics at the individual level. Because personality arise from stable variance components, applying this framework in behavioural research improves our ability to estimate the underlying behavioural phenotype upon which environment and selection can act. Conceptually, animal personality is commonly partitioned into group mean as well as among- and within-individual variance (Barbosa & Morrissey, 2021; Royauté & Dochtermann, 2021). Among-individual variance captures consistent differences between individual behavioural intercepts and therefore represents the component of variation underlying personality, whereas within-individual variance describes the extent to which an individual behaviour fluctuates around its own mean over time. Other sources of behavioural variation, such as behavioural plasticity can reflect predictable modulation of behaviour in response to changing environmental conditions, while repeatability provides the statistical framework for estimating the proportion of total variance attributable to among-individual differences and importantly, reflects the temporal and contextual consistency of a behavioural trait (Dochtermann et al., 2014; Wilson, 2018).

Understanding the structure of behavioural variation is particularly relevant for species kept under captive conditions. Behavioural traits influence how individuals interact with their environments, including access to resource, social interactions, and exposure to environmental challenges (Huntingford et al., 2006; Prentice et al., 2022). Therefore, partitioning behavioural variance can provide valuable insights beyond average behavioural responses by revealing the underlying behavioural architecture. For instance, differences in mean responses may suggest alternative effects between groups, but such comparisons alone provide limited insight into the underlying behavioural structure. An animal personality framework enables a deeper investigation by examining how among⍰individual variation vary between groups, and by assessing the relative contributions of genetic and environmental factors (Prentice et al., 2022; Richter & Hintze, 2019). Behavioural traits that align in response to rearing environmental may cope more effectively, whereas mismatches between behavioural traits and environmental conditions can lead to chronic stress and ultimately compromised welfare. Therefore, integrating animal personality research provides a valuable framework to diagnose welfare in hatchery systems (Huntingford, 2004, 2020; Huntingford et al., 2006).

Behavioural trait such as boldness may be closely linked to functional phenotypes, for instance bold individuals typically take greater foraging risks, increasing food intake and potentially accelerating growth (Biro & Post, 2008; Biro & Stamps, 2008). These behaviours may also be linked to metabolic processes, as bolder and more active fish generally exhibit higher metabolic demands, which can influence energy allocation to growth and neural development (Careau et al., 2003; Careau & Garland, 2012; Réale et al., 2010). Such association between behaviour and other traits reflect the broader concept of animal personality, in which a behavioural trait can covary and/or correlate with a trait to support adaptive responses to environmental conditions often know as coping-style (Dammhahn et al., 2018; Royauté et al., 2018). Although these relationships are both species- and context-dependent, they are consistently shaped by environmental conditions (Sih et al., 2015; Wolf et al., 2007). In hatcheries, for example, the expression of a behavioural trait is strongly influenced by environmental complexity, in which complex environments tend to promote species-specific behaviours and reduce maladaptive response, whereas plain conditions can restrict nature-based behavioural expression (Brydges & Braithwaite, 2009; Huntingford et al., 2006). Yet, responses to environmental complexity remain context specific and can be contradictory, ranging from decreasing activity and aggression in brown trout (*Salmo trutta*) (Hojesjo et al., 2004; Watz et al., 2019) to increasing territoriality and enhanced cognitive performance in Atlantic salmon (*Salmo salar*) (Prentice et al., 2025; Rosengren et al., 2017). But while environmental complexity affects behavioural traits it can also affect functional trait such as growth rate and brain sizes, suggesting that behaviour, growth, and neural traits may emerge through coordinated multivariate phenotypic plasticity (Ebbesson & Braithwaite, 2012; Jonsson & Jonsson, 2014; Näslund & Johnsson, 2016; Shumway, 2008).

Because environmental complexity can simultaneously influence behaviour, growth, and brain development, these traits are unlikely to respond independently. Instead, they may shift together through coordinated phenotypic adjustments, underscoring the limitations of single-trait perspectives. The classic view of phenotypic plasticity typically focuses on a single trait exposed to contrasting experimental conditions (e.g., feeding modalities, temperature gradients, or environmental complexity), with attention placed on a focal phenotype such as morphology, physiology, growth or behaviour. This approach often overlooks the insights gained from examining how multiple phenotypic traits respond jointly and influence one another (Metcalfe, 2024; Metcalfe & Monaghan, 2003; Nielsen & Papaj, 2022; Nussey et al., 2007). For instance, shifts in behavioural responses to benthic prey can increase growth rates in Arctic charr (Horta-Lacueva et al., 2023; Kristjánsson et al., 2018), as foraging behaviours that enhance access to benthic resources increase food intake and consequently promote growth. Because phenotypic plasticity operates across interconnected trait networks, such behavioural shifts may simultaneously influence personality, growth rate, and brain size. A multivariate perspective on phenotypic plasticity is therefore particularly well suited for examining how environmental complexity shapes coordinated phenotypic responses.

The Arctic charr is a salmonid species endemic of alpine and subarctic regions. It is an excellent model for studying phenotypic plasticity due to its genomic capacity to generate a wide range of alternative phenotypes in response to contrasting environmental conditions (Adams et al., 2003, 2007). While several studies have documented the existence of consistent among-individual variation in behavioural trait in Arctic charr (Benhaïm et al., 2023; Dellinger et al., 2023; Ladurée et al., 2026; Philip et al., 2022), no research has yet investigated how environmental complexity influences the long-term consistency of among-individual variation in behaviour and associated phenotypic traits. In fish species personality is often assessed over short time scales (days to weeks); however, such assessments may confound plastic responses with genetic or developmental effects. Among⍰individual behavioural variation may appear repeatable simply because individuals are measured in a consistent environment, potentially obscuring variation arising from genetic or developmental sources (Niemelä & Dingemanse, 2017; Wilson, 2018; Zsebők et al., 2017). Consequently, assessing personality over longer time periods relative to the species’ lifespan may be more informative (Dingemanse & Wright, 2020). Addressing this gap is particularly relevant for understanding the intrinsic character of behaviour as a phenotype.

Here we examined how structural environmental complexity shapes personality in Arctic charr and their co-varying functional phenotypes, specifically growth rate and brain size. We propose that environmental complexity promotes alternative behavioural and functional phenotypes through multivariate phenotypic plasticity. We predicted that fish exposed to complex environments would exhibit lower mean boldness and reduced within- and among-individual behavioural variation. Because environmental complexity often reduces swimming activity and may free energetic resources for somatic and neural development, we also expected enriched conditions to increase growth rates and enlarge or differentiate brain regions. Finally, we predicted that the relationships among boldness, growth, and brain size would differ between rearing environments, generating context-dependent trait associations consistent with multivariate phenotypic plasticity.

## Material and Methods

### Arctic charr Collection and Husbandry

Arctic charr from the breeding station of the University of Iceland at Hólar (Hólar, Iceland) were used in this experiment. A single male was crossed with three females, and the resulting half-sib families were mixed before eggs were incubated at 04.00 °C. After hatching, the batch was split into six 20.00-litre-tanks with a biomass of 13.80 g per tank (644.00 g.m^-3^), with three replicates assigned to each treatment. After the onset of first exogenous feeding (73 days post-hatching; thereafter dph), Arctic charr were exposed to structural environmental complexity, generated in three dimensions: vertically with a green plastic plant, and horizontally with five black pebbles. The second triplicate, representing the plain treatment, was kept bare as white background tanks. Water temperature was maintained at 5 ± 1°C throughout the experiment. Flow rates increased from 48 l. hour^-1^ at hatching to 120 l. hour^-1^ to maintain 100% oxygen saturation on a 12:12□h light cycle. Fish were fed commercial pellets three times daily (9:00am, 1:00pm, 4:00pm). Total Ammonia Nitrogen remained undetectable throughout the experiment.

## Experimental Design

### Sampling Desing

A randomly selected sample of 32 individuals (192 fish in total across 6 tanks) were tagged. Tagging was performed at 259 dph under anaesthesia (2-phenoxyethanol) at a concentration of 310 ppm. A small incision was made between the pectoral fins, and a PIT-tag (Passive Integrated Transponder 1.4 x 8 mm FDX-B PIT tags, Oregon RFID EU GmbH) was inserted into the peritoneal cavity. Behavioural tests started 170 days after the procedure to ensure full recovery.

### Experimental Timeline

Boldness was tested across 4 repetitions at two temporal scales: a short-term interval of 7 days between repetition 1 and repetition 2, followed by a long-term interval of 130 days between repetition 2 second and repetition 3, and finally another 7 day-interval, between repetition 3 and repetition 4. Body mass (g) and fork length (cm) were measured after each behavioural trial. Individuals were euthanized at 470 dph and brains were collected, fixed, and preserved for later measurement (Figure 1).

**Figure 1:**
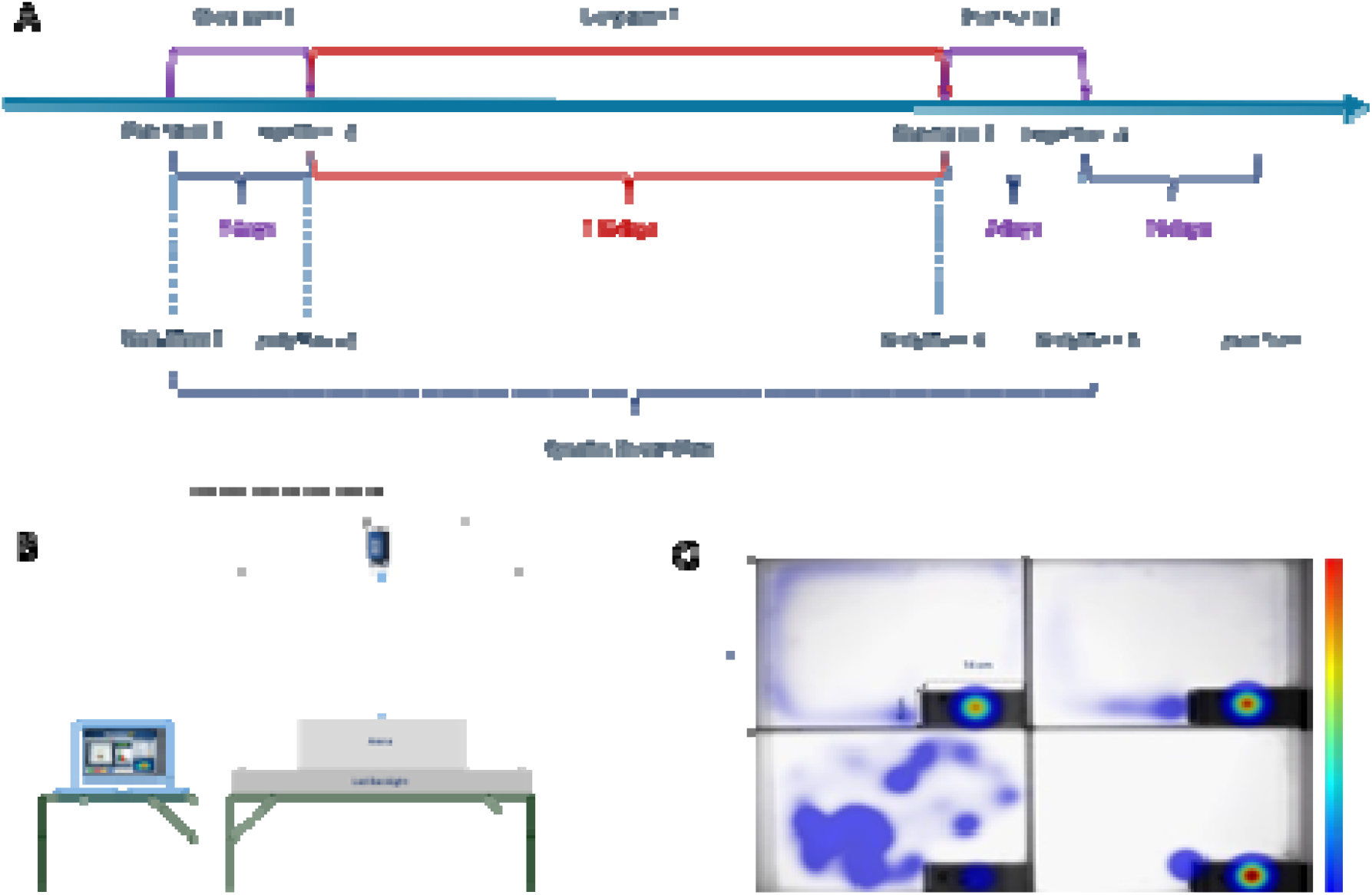
Experimental design and behavioural trait measure setup. A. Experimental design timeline. B. Behavioural trait recording setup. C. Open field with Shelter Arena.

## Behavioural Trait

### Boldness Test: Open Field Test with Shelter

Boldness was measured in an open field arena (30 cm x 40 cm) with a shelter (6 cm x 14 cm – with a roof and front hatch) positioned in the left corner on a white LED backlight (110 x 110 cm, Noldus, the Netherlands). A camera (Basler Ace acA1920-150 mm camera Germany, 30 fps) was mounted above (112 cm) the arenas and connected to a computer for video capture. Trials were recorded using Ethovision XT15 tracking software (Noldus, The Nertherlands). The individual was introduced into the shelter through the roof hatch. After a 5 minutes acclimation period in the shelter, the front hatch was lifted, allowing the individual to explore the open arena for 20 minutes. The decision to exit the shelter into the open area was entirely voluntary as the arena was designed as a non-forced test, allowing individuals to stay hidden in the shelter. After 20 minutes, individuals were placed in a separate aerated container of freshwater and an anaesthetic of 2-phenoxyethanol at a concentration of 310 ppm was used to allow accurate recording of body mass (g) and fork length (cm), after which the individuals were returned to their home tanks upon recovery. Acclimation and trial durations were based on previous studies conducted on the same species (Benhaïm et al., 2023; Dellinger et al., 2023; Philip et al., 2022).

### Behavioural Data Generation

Using Ethovision XT15 tracking software (Noldus, The Nertherlands), the arena was virtually divided into the following zones: entry, border, and centre. For each individual, the following variables of interest were extracted: (1) the total distance moved (i.e., the distance travelled by the centre point of the individual (cm)), (2) the latency time to emerge from the shelter in second (s), (3) the total time spent in the shelter, the centre zone, the border zone and the entry zone (s), (4) the mean velocity (i.e., the distance moved by the centre point expressed in body length per second (body length.s^-1^)), and (5) the absolute angular velocity (i.e., expressed in degrees per second (° s^−1^ )).

### Growth Rate

To track individual growth trajectories, the Specific Growth Rate (SGR) was calculated between repetition 1 and 4 using Equation 1 where *m*_*i*_ and *m*_*f*_ are respectively the initial and the final body length while *Δt* is the time interval in days.

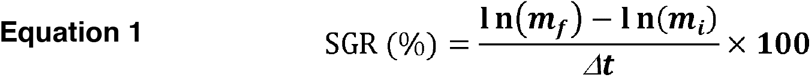

## Brain Size

### Fixation and Conservation

Post euthanasia with anaesthetic overdose, the head of the fish were removed and fixed in 10 % formalin for 21 days, thereafter the head were placed in 1% Phosphate Buffer Saline (PBS) solution.

### Brain Dissection

To extract the whole brain from the cranium, the medulla and olfactory nerves were severed by incision behind the cerebellum as well as in front of the olfactory bulbs, respectively. Brains were placed on a petri dish containing methylcellulose gel (5% concentration) to keep it stable during imaging. Photographs were taken using a fixed Canon 7D camera fitted with a 50mm lens, illuminated by two parallel neon lights (8W, 4000K, 800 lm, spaced 40 cm apart) and placed 15 cm above the brain. Brains were photographed from both dorsal and right lateral views.

### Brain Size Data Generation

Brain volume was quantified using ImageJ. The length, width and height of four neural regions (i.e. olfactory bulbs, telencephalon, optic tectum and cerebellum – Figure 2) were measured and used to calculate regional volumes with an ellipsoid formula where *V* is the estimated volume of the brain region, Length the maximum anterior–posterior dimension, Width the maximum medial–lateral dimension, Height the maximum dorsal–ventral dimension and *π/6* the geometric constant converting a rectangular prism to an ellipsoid approximation (Equation 2) (Peris Tamayo et al., 2020).

**Figure 2:**
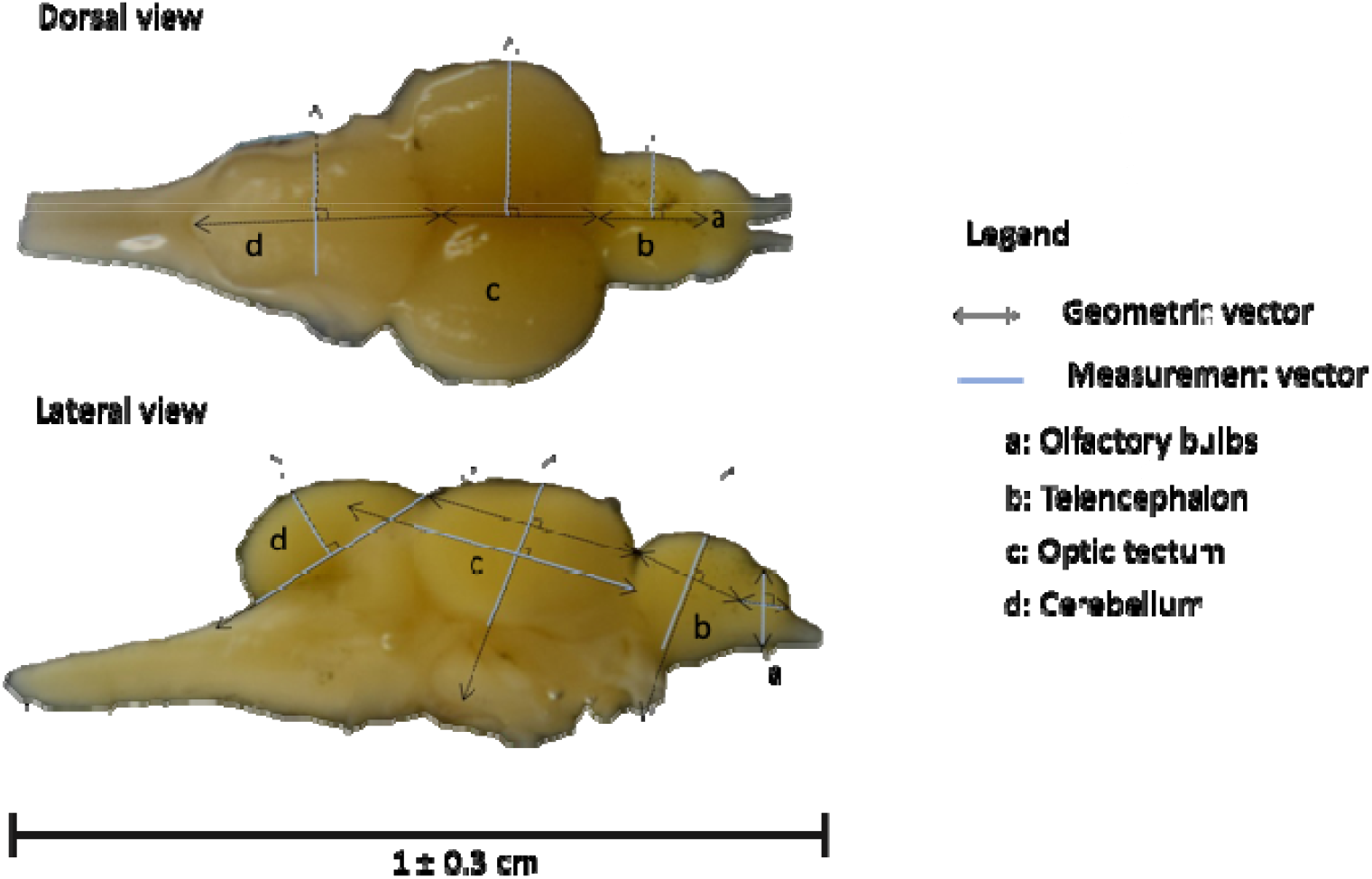
Diagram of the brain regions (a. Olfactory bulbs, b. Telencephalon, c. Optic tectum, d. Cerebellum) and measurements vectors.

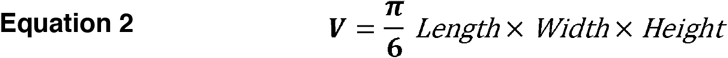

## Statistical Analysis

### Boldness

#### Boldness Score Calculation

The boldness score was quantified using a principal component analysis (PCA) on the variables recorded during the behavioural data generation: total distance moved, velocity, duration spent in the centre of the arena, border zone duration, total time in shelter, latency to exit the shelter, and absolute angular velocity. Variables were centred and scaled (mean = 0, SD = 1) prior to analysis. The PCA was performed using data from the first behavioural trial (Repetition 1) to define a common behavioural axis. Scores from subsequent trials (Repetitions 2, 3 and 4) were projected onto the PCA space using the *predict* function, ensuring that all trials were evaluated against the same boldness structure and avoiding pseudo-repeatability. The first principal component (PC1 (60.50%)) was used as the boldness score, representing a continuum from highly active and exploratory individuals (high PC1 values) to cautious individuals showing greater shelter use (low PC1 values).

#### Behavioural Trait Computation

The variance components of the boldness were computed using a Bayesian linear mixed-effects model implemented in MCMCglmm (Hadfield, 2010). In order to compute a single model and address different timescales between two different treatments the dataset was restructured. A new variable named Serie was created.

This variable groups trial repetitions: repetitions 1 and 2 were assigned to Serie A (Short-term 1), while repetitions 3 and 4 were assigned to Serie B (Short-term 2). Finally, it generates a new identifier called ID_Short_Term by combining each individual ID with its corresponding, allowing individuals to be tracked separately across the two short⍰term behavioural sessions.

A non-informative Inverse-Wishart (*IW*) prior was computed in Equation 3 with the term *G*^1^ as 2×2 covariance matrix describing how individuals differ in boldness across treatments and how consistent they are between treatments. In this prior, the scale matrix *I*^2^ is the 2×2 identity matrix, representing a neutral starting point that assumes no correlation in boldness between treatments. The degrees-of-freedom parameter was defined as a flat prior, allowing the data to drive the estimates. The term *G*^2^ is a 2×2 covariance matrix and represents short-term repeatability across the two-measurement series (Series A and Series B). The term *G*_3_ is a 1×1 matrix and represented the effect of the tank. As above, the scale matrix *I*_2_ assumes unit variance and zero covariance between the two measures. Finally, *R* represent the residual variance–covariance matrix, capturing the within-individual variance and covariance. Again, *I*_2_ denotes the 2×2 identity matrix, a neutral structure assuming that boldness is uncorrelated and has unit variance across the two treatments so that the prior does not influence the model. The MCMCglmm model was computed as follow in Equation 4.

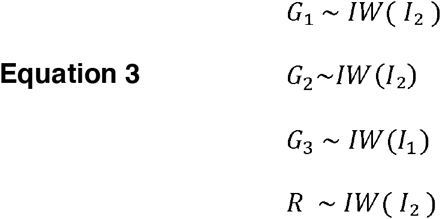

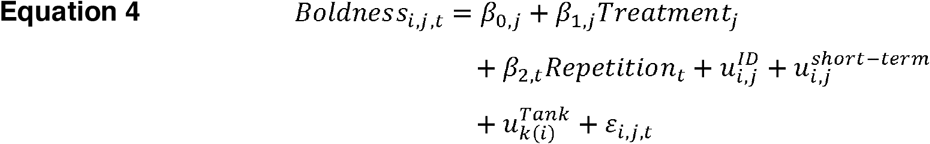

Here *Boldness*_*i,j,t*_ was set as the response variable representing the boldness score of individual *i* in treatment *j* at time *t*. The fixed effects include the independent intercept *β*_0_ *for each treatment j*, the treatment effect *β*_*l,j*_ estimating differences in boldness between treatments, and the repetition effect *β*_*2,t*_ capturing variation across repetition *t*. The model also contains three random-effect components: 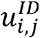, a treatment-specific random effect for individual *i*, this allows to compute the among-individual variance across the long-term period. In order to capture the short-term among-individual variance the treatment-specific for individual *i* is set as a random effect 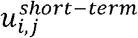. To account for tank, a random effect for individual *i* the experimental tank is set as 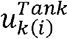. Finally, the residual term *ε*_*i,j,t*_ is estimated separately for each treatment and represents the within-individual residual variation not explained by either short- or long-term individual effects. Together, these components allow the model to partition boldness variation across treatment effects, temporal scales and individual differences. The repeatability (Equation 5) for each temporal scales and treatments are computed using the variances components extracted from the main model and implemented in the repeatability formula below. Where 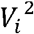 is the square of the among-individual variance by the sum of the among- and within-individual variance 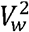 (Houslay & Wilson, 2017; Wilson, 2018).

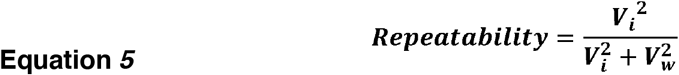

### Trait Covariation and Correlation

The traits components between boldness, growth rate, size of the total brain and described regions, were computed using a Bayesian multivariate linear mixed-effects model implemented in MCMCglmm (Hadfield, 2010). Boldness and the covariate traits were set as two-trait of the multivariate response. Each traits were collapsed into a value by phenotype matrix to allow the integration of the multivariate response in the model. A non-informative Inverse-Wishart (*IW*) prior was computed as shown in Equation 6 with the term *G*_1_ as 4×4 covariance matrix describing how individuals differ in traits (e.g, boldness, growth rate). In this prior, the scale matrix *I*^2^ is the 2×2 identity matrix, representing a neutral starting point that assumes no correlation in boldness between treatments. The degrees-of-freedom parameter was defined as a flat prior, allowing the data to drive the estimates. The term *G*_2_ is a 1×1 matrix and represented the effect of the tank. Finally, *R* represent the residual variance– covariance matrix between the two traits. The MCMCglmm multivariate model was computed as follow in Equation 7. Here Boldness, Trait_*i,p,j*_ was set as the multivariate response variable representing the trait value of individual *i* for phenotype p in treatment *j*. The fixed effects include the independent intercept *β*_*O,p,j*_ estimating differences in traits between treatments. The model also contains two random-effect components: 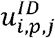, a treatment-specific random effect for individual *i* and phenotype p. To account for tank, a random effect for individual *i* in the experimental tank is set as 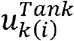 Finally, the residual term *ε*_*i,p,j*_ is estimated separately for each traits.

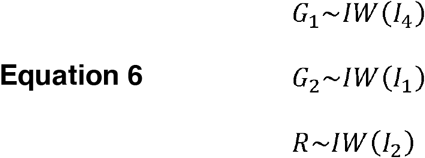

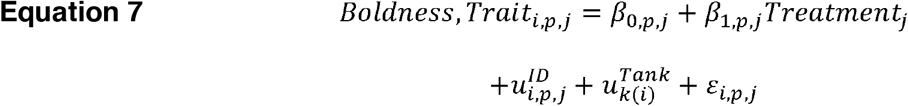

Covariance between traits was estimated separately for each treatment from the posterior samples of the among-individual variance–covariance matrix. The same procedure was applied to obtain the covariance for the complex and plain treatment. Posterior correlations were computed by dividing the covariance between the traits cov_*Boldness,Trait*_ by the square root of the product of the corresponding posterior variances for each trait 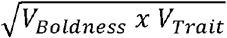 within each treatment (Equation 8).

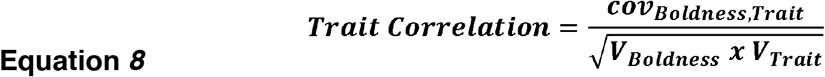

## Results

### Boldness

#### Mean Boldness

Mean boldness did not differ clearly between treatments and across repetitions. Individuals exposed to the plain treatment showed a tendency to be bolder, but the difference was not significant (boldness = 0.290 [-0.119; 0.695], p-value = 0.126) (Figure 3). Mean boldness also showed a temporary decrease at the second repetition (boldness = -0.244 [-0.467; -0.023], p-value = 0.030), but returned to baseline afterwards, with later repetitions not differing from the first (Table 4).

**Figure 3:**
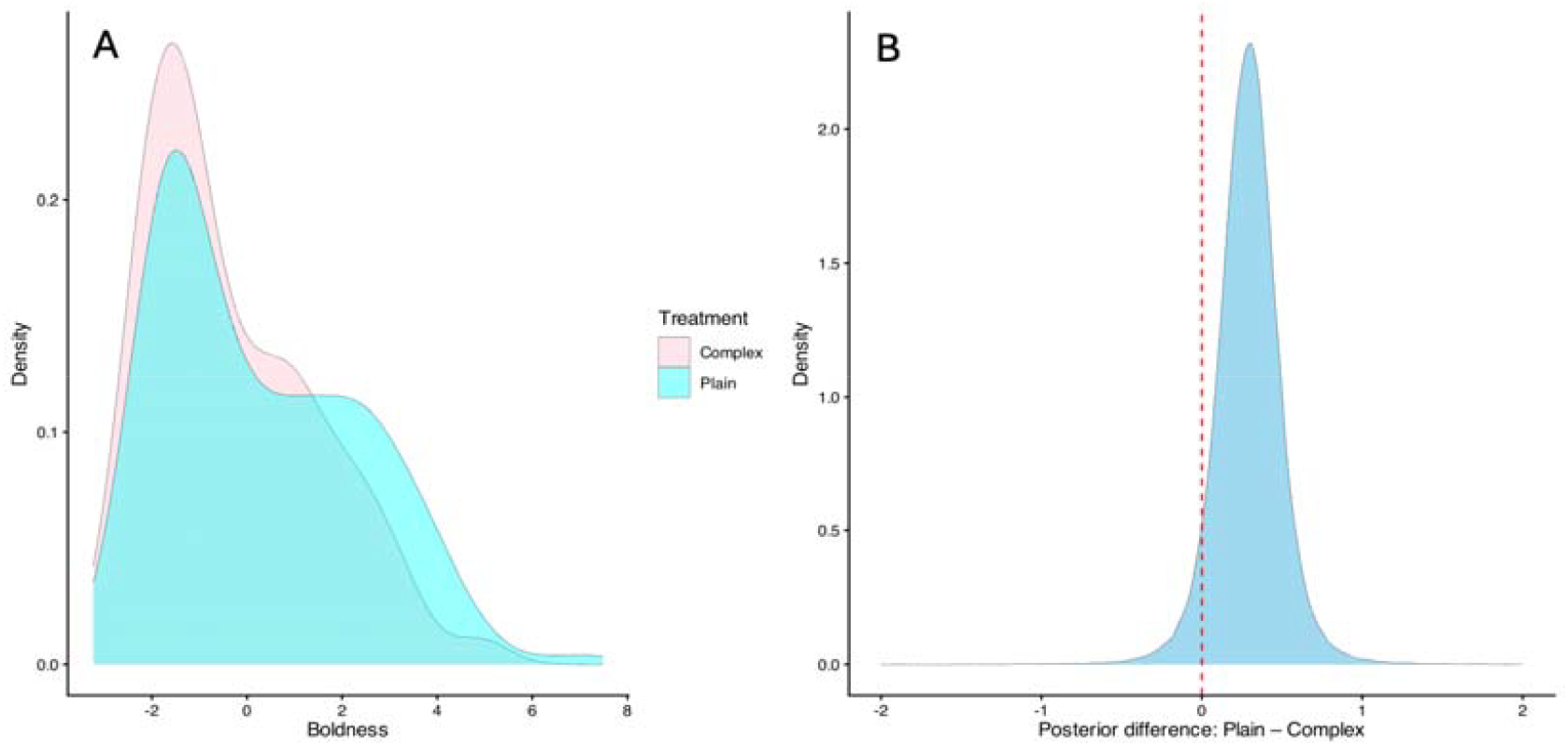
ADistribution of boldness between the plain (cyan) and complex (pink) environment, this figure represents the global distribution of boldness in each treatment, here no differences in mean boldness are observed. B. Posterior difference in mean boldness between the plain and complex treatment. Here the lower confidence interval is crossing the null hypothesis (red dashed vertical line) therefore the difference in boldness between the two treatments is not statistically credible.

#### Among-Individual Variation

Among-individual variance differed across timescales but did not differ credibly between treatments. At the long-term scale, the complex treatment tended to have slightly higher among-individual variance (Vi = 0.221 [0.071; 0.415]) than the plain treatment (Vi = [0.058; 0.205]), but the 95% credible interval for the difference overlapped zero, indicating no strong statistical support for a treatment effect. At the short-term scale, among-individual variance increased in both treatments, with broadly overlapping credible intervals between the complex (0.283 [0.130; 0.488]) and plain (0.301 [0.101; 0.539]) environments (Table 1, Table 5).

**Table 1:**
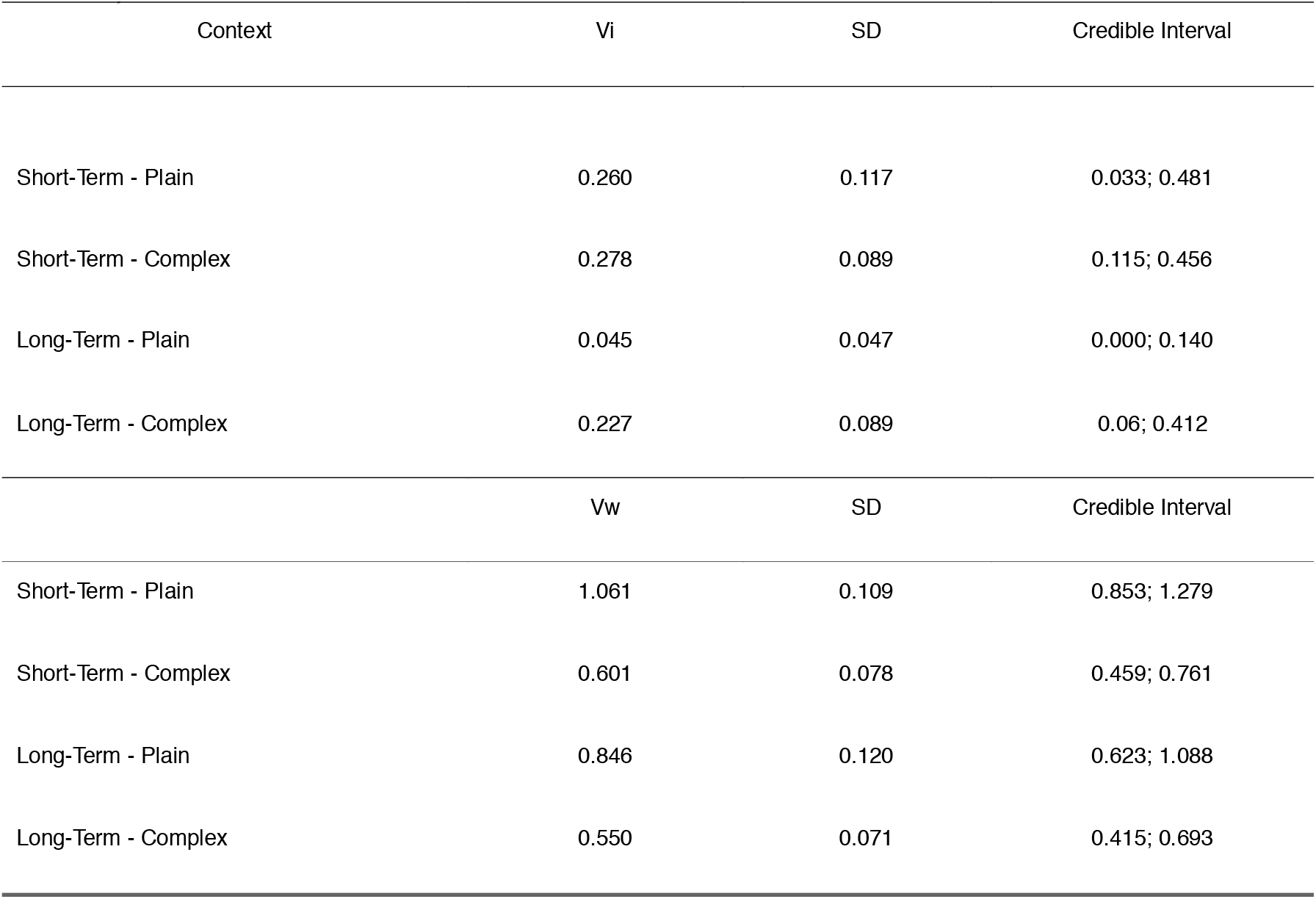
Estimates of among- and within-individual variances (respectively Vi and Vw) with standard deviations and credible intervals across the four context conditions (Short-Term Plain, Short-Term Complex, Long-Term Plain, Long-Term Complex).

**Table 2:** Covariance and correlation matrix of behavioural by phenotypic association. Here for each trait association and each treatment, the covariance as well as the correlation estimates are reported along with their respective confidence interval.

| Trait | Treatment | Covariance | Cov. | Correlation | Corr. |
| --- | --- | --- | --- | --- | --- |
|  |  |  | Credible Interval |  | Credible Interval |
| Boldness x Growth | Complex | -0.003 | -0.09; 0.08 | -0.01 | -0.28; 0.26 |
|  | Plain | -0.003 | -0.07; 0.07 | -0.01 | -0.27; 0.25 |
| Boldness x Brain Size | Complex | -0.080 | -0.31; 0.13 | -0.13 | -0.46; 0.23 |
|  | Plain | 0.053 | -0.25; 0.36 | 0.06 | -0.30; 0.40 |
| Boldness x Olfactory bulbs | Complex | 0.153 | -0.08; 0.44 | 0.22 | -0.14; 0.54 |
|  | Plain | 0.129 | -0.17; 0.46 | 0.15 | -0.21; 0.49 |
| Boldness x Telencephalon | Complex | 0.017 | -0.23; 0.27 | 0.02 | -0.33; 0.38 |
|  | Plain | 0.233 | -0.03; 0.55 | 0.30 | -0.04; 0.60 |
| Boldness x Optic tectum | Complex | -0.059 | -0.30; 0.15 | -0.10 | -0.44; 0.25 |
|  | Plain | 0.051 | -0.27; 0.36 | 0.06 | 0.31; 0.39 |
| Boldness x Cerebellum | Complex | -0.18 | -0.52; 0.10 | -0.23 | -0.56; 0.13 |
|  | Plain | -0.11 | -0.48; 0.24 | -0.11 | -0.46; 0.24 |

#### Within-Individual Variation

Within-individual variance differed between treatments and across temporal scales. At the long-term scale, individuals in the plain treatment exhibited higher withinindividual variance (Vw = 0.83 [0.65; 1.09]) compared with those in the complex treatment (Vw = 0.54 [0.41; 0.70]). When short-term individual variance was included, total within-individual variance increased in both treatments but remained greater in the plain (Vw=1.08 [0.88; 1.31]) than in the complex (Vw=0.60 [0.47; 0.79]). These patterns indicate that Arctic charr in the plain treatment showed greater within-individual behavioural variance across observations. In contrast, individuals in the complex treatment showed greater stability in their behaviour, explained by lower estimates of within-individual variance both at short- and long-term scales (Table 1, Table 5).

#### Repeatability

Repeatability estimates, effectively reflecting the personality structure of the individuals in this study, differed between treatment and across temporal scales. Short-term (7 days) repeatability was not different (posterior mean = 0.077 [-0.133– 0.308]) between plain ((R = 0.262 [0.093; 0.427]) and complex (R = 0.338 [0.175; 0.505]). Here it is nevertheless notable that early sign of unstructured personality appears in the plain treatment with the very weak lower confidence interval (0.093) (Figure 4, Table 3).

**Table 3:** Repeatability estimates for each time periods and treatments. Values include standard deviation (SD) and 95% credible intervals.

| Context | Repeatability | SD | Credible Interval |
| --- | --- | --- | --- |
| Short-Term - Plain | 0.233 | 0.096 | 0.044; 0.412 |
| Short-Term - Complex | 0.332 | 0.082 | 0.171; 0.49 |
| Long-Term - Plain | 0.048 | 0.048 | 0; 0.146 |
| Long-Term - Complex | 0.286 | 0.088 | 0.111; 0.456 |

**Table 4:** Posterior estimates from the Bayesian model assessing the effects of treatment and trial repetition on behavioural responses. Values include posterior means, 95% credible intervals, and associated p-values.

| Effect | Posterior Mean | Credible Interval | p-value |
| --- | --- | --- | --- |
| Intercept | -0.17304 | -0.494; 0.154 | 0.2499 |
| Treatment (Plain) | 0.29026 | -0.119; 0.695 | 0.1266 |
| Repetition 2 | -0.24469 | -0.467; -0.023 | 0.0302* |
| Repetition 3 | 0.11386 | 0.122; 0.347 | 0.3416 |
| Repetition 4 | 0.13989 | -0.086; 0.382 | 0.2418 |

**Table 5:** Posterior estimates comparing short-term and long-term differences between complex and plain treatment for the following variance components: repeatability, among- and within-individual variance (Vi and Vw). Values include posterior means, standard deviations (SD) and 95% credible intervals.

| Comparison | Repeatability Posterior | SD | Credible Interval |
| --- | --- | --- | --- |
|  | Mean |  |  |
| Short-term (Complex – Plain) | -0.099 | 0.126 | -0.342; 0.149 |
| Long-term (Complex – Plain) | -0.237 | 0.098 | -0.423; -0.032 |
| | $V_i$ Posterior Mean | SD | Credible Interval |
| Short-term (Complex – Plain) | 0.018 | 0.147 | -0.004; 0.393 |
| Long-term (Complex – Plain) | 0.182 | 0.100 | -0.277; 0.303 |
| | $V_w$ Posterior Mean | SD | Credible Interval |
| Short-term (Complex – Plain) | -0.460 | 0.134 | -0.730; -0.203 |
| Long-term (Complex – Plain) | -0.296 | 0.141 | -0.585; -0.036 |

**Figure 4:**
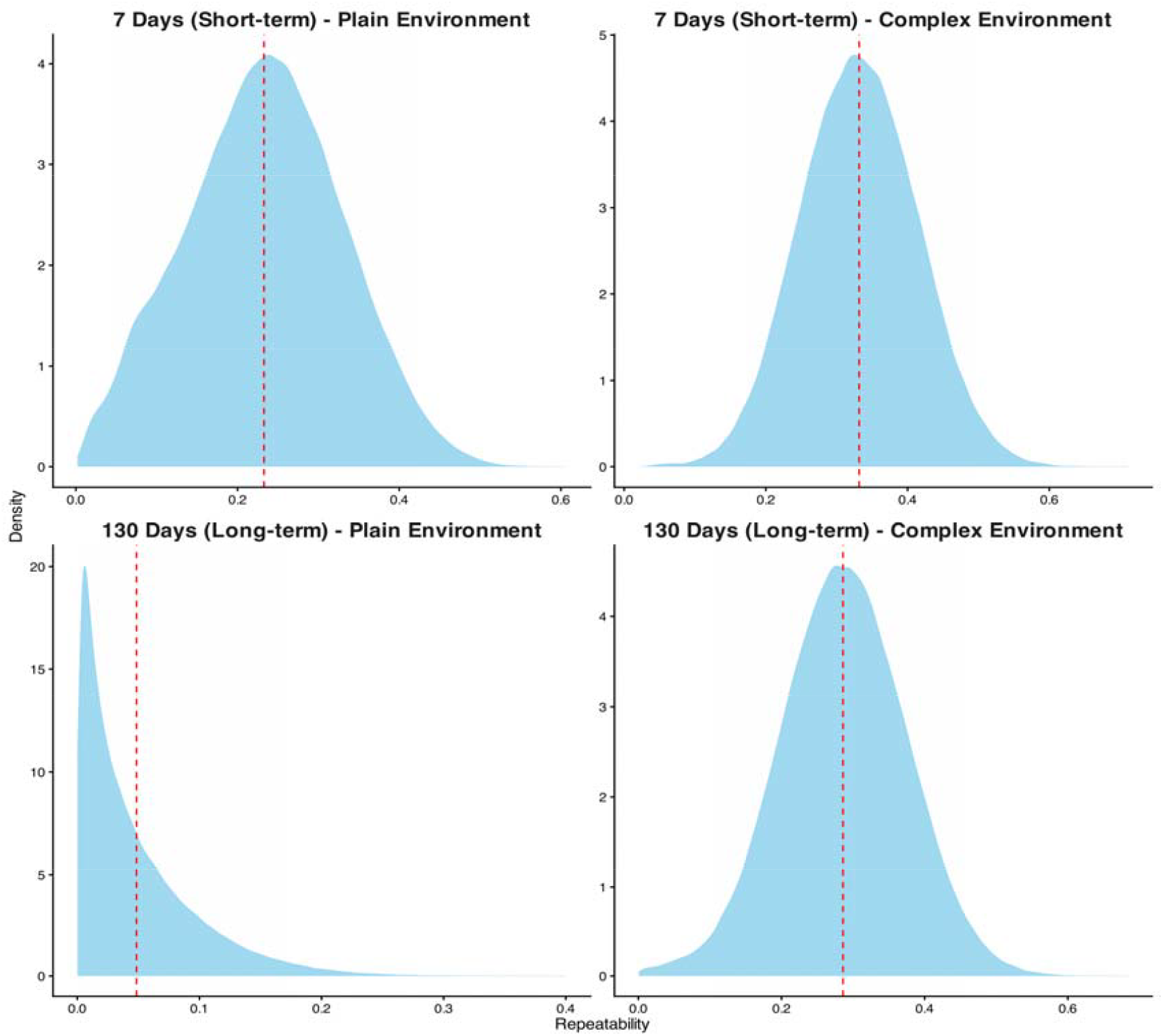
Repeatability estimates for boldness between plain and complex environments and across short and long-term time periods.

On the long-term time scale, the posterior difference between the treatment is significant (posterior mean = 0.213, [0.020; 0.388]), indicating difference in repeatability. Here, repeatability estimates collapse in the plain treatment (R = 0.051 [0.004; 0.171]) and remain substantially higher in the complex treatment despite a decrease in the lower confidence interval (R = 0.263 [0.089; 0.431]) (Figure 5, Table 5).

**Figure 5:**
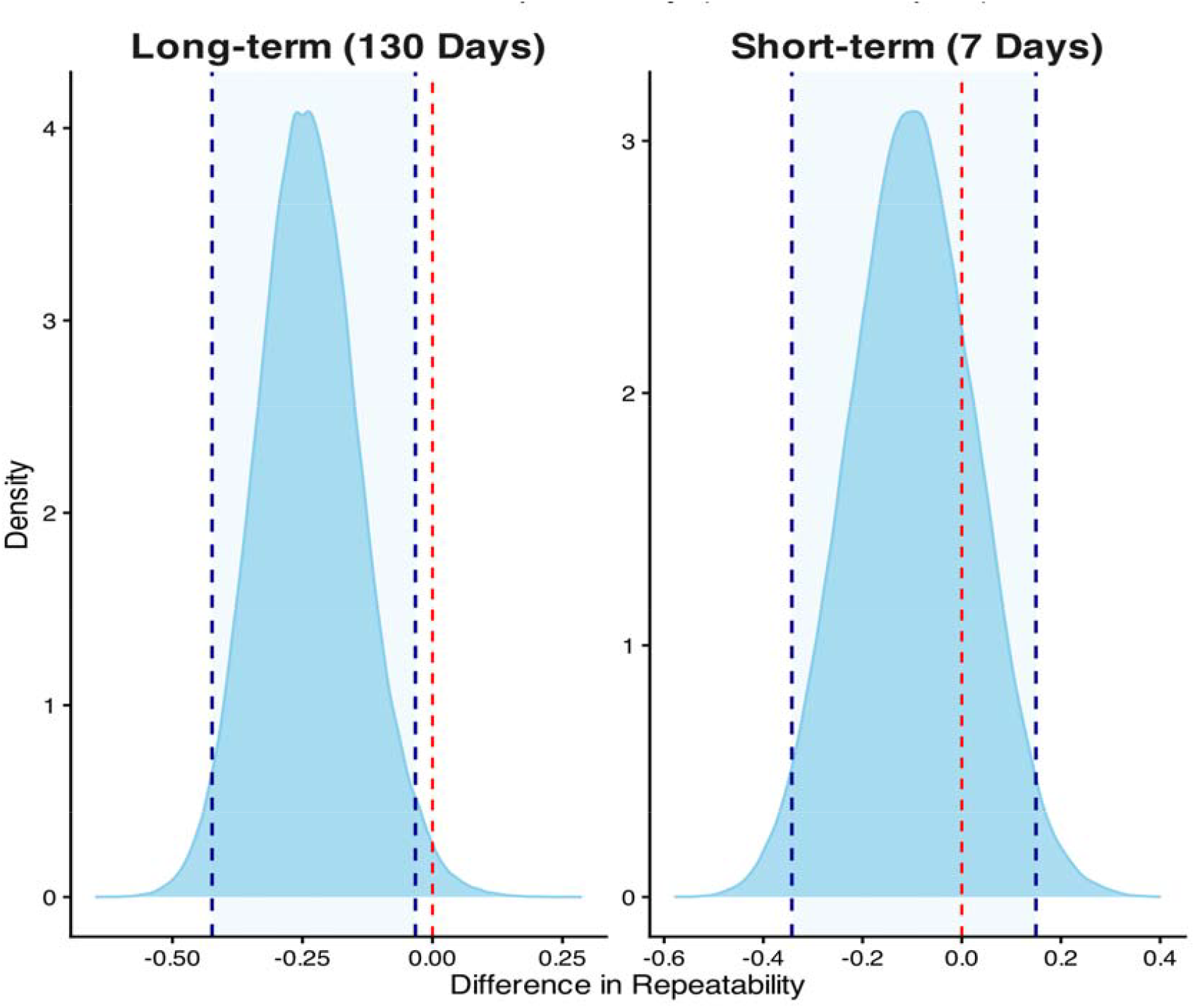
Posterior difference in repeatability estimates between the plain and complex treatment and time periods. Here differences in repeatability are significant over the long-term period. Over the shortterm period the lower confidence interval is crossing the zero (red dashed vertical line) therefore the difference in boldness between the two treatments is not statistically credible.

### Trait Covariation and Correlation

Neither the growth rate (posterior mean=0.59□[–0.48;□1.64]) nor the size of the total brain or individual regions (posterior mean=0.59□[–0.63;□1.85]) were credibly affected by the treatments. In addition, no covariation or credible correlation was found between boldness and the specific brain regions. It is nevertheless worth noting that the estimated correlation between boldness and telencephalon size tended to be positive in the plain treatment (r=0.23□[–0.03;□0.55]). However, the lower bound of the credible interval includes zero, indicating that the evidence for a positive relationship remains weak (Table 2).

## Discussion

This study aimed to investigate how environmental complexity shapes personality in Arctic charr. In addition, we evaluated whether boldness covaries with key functional phenotypes such as growth rate and brain size within a multivariate phenotypic plasticity framework. We have demonstrated that environmental complexity can to some extent maintain personality structure. Although mean boldness did not differ between plain and complex treatments, individuals exposed to complexity exhibited long-term consistency, with higher repeatability estimates associated with reduced within-individual variation. These patterns suggest that environmental complexity does not simply shift the mean level of a behavioural trait but rather reshapes how the trait is expressed. Furthermore, boldness neither covaried with nor was correlated with growth rate or brain size.

### Arctic charr personality and environmental complexity

The mean value of boldness was not significantly higher in the plain treatment, suggesting that exposure to either a plain or complex environment does not modulate average levels of boldness in this species. This contrasts with previous findings reporting higher boldness in plain environment, notably in Atlantic salmon (Roberts et al., 2011; Church et al., 2018). However, the present findings align with multiple other studies showing that structural environment does not necessarily determine average boldness levels, including work on the same strain of Arctic charr, as well as studies in zebrafish and lizard (DePasquale et al., 2016; Philip et al., 2022; De Meester et al., 2022).

We expected that, mechanistically, the absence of refuge in the plain treatment may compel individuals to adjust their behaviour to cope with environmental constraints through behavioural compensation (C. Adams & Huntingford, 2005; Näslund & Johnsson, 2016). For instance, an individual that remains inactive or overly cautious in an environment with limited hiding opportunities may suffer reduced foraging success. Such adjustments reflect behavioural modulation, whereby individuals must frequently modify their behaviour in response to situational demands (Conrad et al., 2011; Cutts et al., 2001; Sih et al., 2004). In our study, this interpretation is supported by the lack of long-term repeatability, associated with high within-individual variation in boldness observed in the plain environment.

Nevertheless, short-term repeatability estimates did not differ between treatments, and Arctic charr in the plain treatment exhibited consistent among-individual variation over a 7-day period. However, after 130 days, repeatability drastically decreased in the plain treatment, while remaining stable in the complex. This does not imply that Arctic charr exposed to plain environments fail to express boldness. Rather, repeatability represents an upper bound of heritability, including both genetic and permanent environmental effects (Barbosa & Morrissey, 2021; Dochtermann et al., 2019); therefore, boldness in both treatments may still own a genetic basis, but the complex treatment appears to foster more stable and consistent among-individual variation, effectively buffering personality expression. In contrast, the plain treatment may induce more context-dependent behavioural responses to cope with stressful environmental conditions. In other words, behaviour may be driven more by external constraints than by intrinsic behaviour. This phenomenon, often referred to as personality suppression, has been demonstrated in three-spined stickleback, where consistent boldness differences diminished following changes in foraging context, indicating that environmentally induced stressors can override the expression of inherent behavioural traits (MacGregor et al., 2021; McDonald et al., 2016).

### Arctic charr Personality and Brain size

The absence of an overall effect of environment on total brain and on specific brain regions, together with the absence of a statistically supported covariation between boldness and telencephalon, suggests that the cognitive demands imposed by the environmental treatments were likely insufficient to drive broad neuroanatomical divergence. In the plain treatment, we observed a trend toward a positive correlation between boldness and telencephalon. Thus, we could argue that such feedback may be present, although they may be significantly detectable with greater power, this pattern does not align with theoretical expectations that personality and brain size can coevolve in coordinated ways when personality is unstructured. The concept of boldness–brain covariation proposes that personality is linked to predictable patterns of neural allocation. Bolder individuals often rely on rapid, proactive decision⍰making strategies, reducing the need for extensive sensory processing, whereas shyer individuals tend to invest more heavily in neural circuits associated with vigilance, risk assessment, and social monitoring (Réale et al., 2010; Sih & Del Giudice, 2012). Accordingly, this framework predicts that behavioural relationships between behaviour and brain size should be more apparent at the level of specific brain regions than in overall brain size. In fishes, several studies have shown that bolder individuals tend to have smaller or differently shaped brain regions associated with sensory integration or social processing, (Gonda et al., 2013; Kotrschal et al., 2014). Thus, the trend observed between boldness and telencephalon size in the plain treatment is not consistent with these theoretical expectations because personality structure collapse on the long term.

### Arctic charr Personality and Growth Rate

Despite theoretical predictions that boldness and growth rate should covary, we found no evidence supporting this relationship in our study. This lack of covariation contrasts with predictions derived from pace⍰of⍰life and behavioural–metabolic syndrome frameworks, which propose that bolder, more proactive individuals should exhibit faster growth due to higher foraging motivation and greater boldness (Réale et al., 2010; Biro & Stamps, 2008). Instead, growth rate appeared neutral and did not covary nor correlate with personality. This absence of correlation suggests that there is no indirect selection for bolder phenotypes contrarily to observations in other fish species (Lu et al., 2022). This result may be an artefact from the common garden experiment due to the same amount of food distributed equally to both treatments. Several studies have reported no effect of environmental complexity on growth rate, with contrasting findings across studies suggesting that the influence of environmental enrichment on growth is largely species-specific and reflects the ecological characteristics of each species (Arechavala-Lopez et al., 2019; Arechavala□Lopez et al., 2022; Gesto & Jokumsen, 2022).

### Animal Personality and Welfare

Our findings have clear implications for fish welfare through the lens of nature-based welfare, as proposed by Huntingford (2006). This framework emphasises that captive environments should enable animals to express the full range of behaviours they would naturally perform in the wild, rather than constraining them to a narrow set of responses imposed by environmental limitations. In the present study, fish held in the plain environment showed unstable expression of boldness through time. It is argued that temporal environmental variation may result in higher within individual variation in behaviour (Stamps and Groothuis, 2010). This concurs with our study, where fish exposed to the plain treatment, are also potentially more exposed to punctual external disturbance in absence of physical shelters, exhibited higher within individual variation compared to fish raised in the complex treatment.

If the observed higher within individual variation results from a higher exposure to various and unpredicted disturbances, fish may also experience altered stress response. Such behavioural modulation responses suggest that the plain environment may not support the expression of natural behavioural repertoires, thereby reducing welfare standard. Chronic stress exposure is a well-recognised source of welfare issues in captive animal populations, often leading to behavioural alterations and plain environments may exacerbate this by exposing individuals to unpredictable stimuli. In contrast, the complex environment, by providing shelter, landmarks, and opportunities for context-appropriate behavioural choices closely aligns with nature-based welfare.

## Conclusion

This study demonstrates that repeatability in boldness is consistent over 130 days in individuals exposed to a complex treatment, whereas it decreases drastically after 7 days in individuals exposed to the plain, supporting the phenomenon of personality suppression. Here it is important to note that our design is not suitable to estimate additive genetic variance through narrow-sense heritability of boldness and we do recommend future study to address this gap to complement the interpretation of variance partitioning in animal welfare. We also found no effect of environmental complexity on mean growth rate or brain size, nor any evidence of covariation or correlation between these traits and boldness. Although this does not rule out the existence of behaviour-brain associations in Arctic charr, the metrics used in the present study may not have been sufficiently sensitive to detect such relationships. We therefore recommend that future studies investigate more refined neuroanatomical measures, such as neuronal density rather than brain ellipsoid volume, or directly quantify cognitive traits within a quantitative personality-cognition behavioural syndrome framework. Together, our findings provide compelling evidence that partitioning behavioural variance can reveal subtle welfare-related effects in groups exposed to contrasting environmental conditions. Consequently, we propose that the animal personality framework represents a valuable tool for addressing key challenges in animal welfare assessment and diagnosis.

## Acknowledgements

Joris Philip, Marion Dellinger and David Benhaïm were supported by the IRF grant number no. 195615-051. At the time of writing this manuscript, Joris Philip was supported by an independent scholarship from the Fisheries Society of the British Isles, hosted at the University of Glasgow. Gabrielle Ladurée, Audrey Prat and Sabine Lobligeois were supported by the aquaculture and fish biology department of University of Iceland and Hólar University. The authors are very grateful to Amber Monroe (Hólar University) for technical support to the Arctic charr husbandry. We thank the Hólar University breeding station for proving the Arctic charr subject to this study. We also thanks Dr. Elizabeth Mittell (University of Edinburgh), Prof. Colin Adams (University of Glasgow) and Dr. Marie-Laure Bégout (Ifremer, France) for early revision and comments on the manuscript. Finally, we would like to thank the member of the aquaculture and fish biology department of University of Iceland and Hólar University for supporting this research.

## Authors Contribution

Joris Philip: Original Idea, Conceptualisation, Data collection, Data generation, Data Analysis, writing original manuscript, writing and editing. Gabrielle Ladurée: Data analysis, writing and editing. Audrey Prat: Data collection. Marion Dellinger: Data collection, writing and editing. Sabine Lobligeois: Conceptualisation, writing and editing. David Benhaïm: Funding acquisition, conceptualisation, writing and editing.

## Data Availability

Data and code related to this research article can be access through this Github repository https://github.com/JorisPhilipResearch/MS007.

